# Epigenetic Regulation Explains The Functionality Behind Colon Cancer Specific Biomarker Septin9

**DOI:** 10.1101/2023.01.31.526413

**Authors:** Laura Vizkeleti, Csaba Kiss, Viktoria Tisza, Aniko Szigeti, Akos Gellert, Istvan Csabai, Lorinc S. Pongor, Sandor Spisak

## Abstract

Despite advancements in early cancer detection and prevention methods, colorectal cancer (CRC) remains a significant global health problem. It is the third most common cancer and the second leading cause of cancer-related deaths worldwide. Additionally, there has been a marked increase of incidence in young adults, and the reasons for this tendency are not fully understood. Therefore, the need for more effective diagnostic methods of assessing disease risk at early stage is crucial.

One of the newly developed blood-based circulating biomarkers with promising potential is the short hypermethylated region located at the Septin9 intronic region. Several clinical studies have proven its performance and applicability. However, the molecular mechanism behind this consistent and recurrent feature present in most of the CRC and related precancerous stages and why it is specific and advantageous for CRC development are poorly understood.

Here, we used comprehensive epigenetic and gene expression profile analyses from different sources of human clinical samples and cell line data to link specific hypermethylation events at the Septin9 intronic loci, which initiate alternative transcription of the Septin9 gene.

Through our investigation of TCGA-COAD RNA-seq samples (n=287), we found that there was no significant difference in global Septin9 levels between normal and tumor samples. However, we did observe a significant alteration in the transcript variant ratio between v1 and v2, suggesting the use of an alternative promoter. Our findings were further supported by our analysis of ATAC-seq data, which revealed that the v2 promoter conferred higher chromatin accessibility, which correlated with the expression of the v2 isoform. However, this was not supported by promoter or enhancer activity as measured by H3K27ac signals. Hypermethylation at the v2 promoter was confirmed in tumor samples, providing a possible explanation for the switch in variants.

Protein sequence analysis confirmed small differences between Septin9 variant ‘A’(v1) and ‘B’(v2). However, AlphaFold2 indicates a substantial difference at the N terminus, which could impact protein phosphorylation. We hypothesize, that variant ‘A’ (v1) and variant ‘B’ (v2) are required for normal cell functions but shifting the balance towards v1 is more favourable for the tumor.

Although very little is known about Septin9 and its function in CRC biology, we are confident that our study will help to emphasize the importance of understanding regulatory mechanisms behind tumor-specific biomarkers and helps to improve the application.

## Background

Based on WHO data, colorectal cancer (CRC) was the third most common cancer type, accounting for 1.93 million new cases (contributing 10.7% of new cases) in 2020, and the second most common cause of cancer related deaths accounting for 916.000 deaths worldwide [1]. In Hungary, CRC has been characterised by increasing incidence and more or less stagnating mortality (~5.000 deaths per year) over the past decade. With 10-11 thousand new cases per year, the incidence of CRC is considerably high in both men and women [2]. Of newly diagnosed patients, ~20% will have metastatic disease at first presentation, and a further ~25% of CRC patients will present metastasis within the next 1-3 years. Approximately 70-75% of patients with metastases are lost within 1 year [3]. Despite recent advances in management of CRC, the usually unresectable metastatic disease still remains challenging [3,4]. Therefore, early detection is essential for successful management of CRC patients. In 2013, Wasserkort et al. defined the methylation pattern of *SEPTIN9* gene on tissues derived from adenoma and colon adenocarcinoma (COAD) separately examining stroma and epithelial abnormalities by bisulfite sequencing of 8 amplicons [5]. They compared these results to both normal- and peritumoral normal tissues. The amplicons tested showed high methylation of Amp4 (in tumor only), Amp5 and Amp6 in both tumor and adenoma epithelium. Based on the human genome assembly hg38, Amp4-6 overlap a CpG island at the *SEPTIN9* locus. This tumor-associated aberrant hypermethylation pattern has already been identified in a high proportion (87%) of early stage (stage I and II) tumors and can also be detected in circulating DNA of the patients’ peripheral blood.

Diagnostic screening for early detection of CRC, adenomatous polyps and precancerous lesions can remarkably reduce cancer-related mortality with up to 60-70% using a recommended starting age of 45-50 years [6–9]. Screening tests can basically be divided into 3 methodological groups: stool-based, endoscopic and molecular tests. Stool-based tests such as faecal occult blood test (FOBT) and multitarget stool DNA test (FIT-DNA) are non-invasive, cost-effective and easily performed approaches. However, in case of FOBT dietary restrictions are needed before testing as it detects heme component of the haemoglobin. Further drawbacks are the high false-negativity rate and poor sensitivity for benign polyps. Although FIT-DNA has a higher sensitivity than FOBT, the false-positivity rate is relatively high (up to 14%), and it is still less sensitive for precancerous lesions with a greater cost [6–9]. Out of many endoscopic tests, colonoscopy is a widely used and preferred screening method of the entire colon due to its high sensitivity, accuracy and possibility to take biopsy from the suspicious lesion. However, a full bowel preparation is needed with a conscious sedation or general anaesthesia causing inconvenience to the patient and possible complications. Many alternative non-invasive procedures are now available that do not require anaesthesia such as flexible sigmoidoscopy (FS), CT Virtual colonoscopy (CT-VC) or Colon capsule endoscopy (CCE). A great advantage of FS is that it is an outpatient procedure without a need for bowel preparation, however, it inspects only the rectosigmoid region. In case of CT-VC and CCE, the main drawbacks are lack of biopsy taking and high cost [6–9]. The third group is the molecular tests based on non-invasive liquid biopsy such as Septin9 test [10]. This qualitative molecular screening kit detects epigenetic changes of Septin9 DNA from peripheral blood using a simple blood draw followed by a methylation specific PCR reaction. Methylation level of Septin9 gene is higher in tumor affected patients, even in early stages, compared to healthy individuals. However, most studies examining the performance of the kit excluded “difficult-to-diagnose” patients and/or did not specify a clear detection threshold. Molecular marker detection, therefore, is still considered an experimental method, a preliminary tool with no clear guideline for their use [6–9]. Its positive result should be verified by colonoscopy or sigmoidoscopy.

Despite the uncertainties of molecular tests of Septin9 or other marker, they can be an important part of diagnostics due to their non-invasive nature. But it is also worth looking carefully at the basic functions of the markers applied. About the higher order organisation of septins itself we know a relatively much from a number of structural studies. To back up the accurate utilization of Septin9 based molecular tests and the interpretation of the resulted outcomes, we aimed to provide an in-depth analysis of Septin9 isoforms and epigenetic regulations.

Septins are evolutionarily highly conserved proteins with slow intrinsic GTPase activity (except for the Septin6 subgroup), which are inherently unstable, forming hexamer or octamer subunits depending on their function. They are also known as the “fourth” component of the cytoskeleton because of their filament-forming property, creating scattered networks in the cytoplasm and cell cortex [11–14]. The dynamics of assembly varies strongly throughout the cell cycle. In terms of their function, they play a role in all processes where membrane rearrangement and/or anchoring is required. Septins are therefore essential for the normal development and function of different organ systems both during and after ontogenesis [15–17]. In the human genome, a total of 13 genes in 4 subfamilies encode more than 30 common or tissue-specific isoforms. The expression level of Septin9 influences the ratio of hexamer-octamer complex, and the composition and expression level of each isoform together determine the higher-order structures of septins [11,15]. Although little is known about the organisation process itself, the biochemical properties and posttranslational modifications such as phosphorylation, acetylation, ubiquitination or sumoylation result in a large number of possible interacting partners [18].

Septin9 localized at 17q25.3 locus is characterized by a complex genomic organization. Besides the relatively high number of coding exons, several alternative transcripts can be found. Some transcripts code the same protein, transcribed from different promoters which have divergent transcription effectiveness [19]. Currently 11 transcript variants are known, encoding 8 protein isoforms. The composition of these isoforms is cell type specific, and fundamentally influences several intracellular processes [20]. Septin9 proteins occur only in stable octamers. Their absence is embryonically lethal, since they are essential for proper mitosis or planal cell polarity during and after embryogenesis [20,21]. They regulate karyokinesis during ana- and telophase of cell division by localizing midbody-exocyst complex, promoting vesicle fusion, which finally leads to the division of daughter cells. Decreased expression of Septin9 results in multinucleated and/or microtubule-bridged daughter cells [22]. Therefore, in general, loss of function of Septin9 leads to genome instability. However, the re-expression of isoforms 1 (A in present nomenclature) or 3 (C in present nomenclature) alone is able to repair this cytokinesis defect, while the presence of isoform 4 (E in present nomenclature), which codes a truncated protein form, has a dominant negative effect. It is assumed that there may be several phosphorylation sites at the N-terminal part of the Septin9 protein essential for its function and are lost with truncation [20]. Despite the possible importance of Septin9 epigenetic alterations in CRC detection, there is a lack of knowledge about the molecular processes that regulate the spatial and temporal expression and isoform composition.

The molecular biology of Septin9 abnormalities is mostly studied in breast and ovarian cancers. Mutation of Septin9 gene is not typical in malignant tumors. In AML (acute myeloid leukemia) and ALL (acute lymphocytic leukemia) the MLL (mixed-lineage leukemia gene)-SEPT9 inframe fusion occurs following a double stranded intronic break in about 10% of *de novo* and therapy resistant leukaemia, [23]. In ovarian cancer overexpression of v1 and v4* (alias v5) variants were detected in serous and mucinous borderline tumors. The hypoxia-induced enhanced translational efficiency of v4* over v4 (alias v6) results in increased i4 (alias ‘E’) protein level, which promotes the malignant and metastatic potential via enhanced motility and resistance to microtubule target drugs (e.g. taxanes) [16,24,25]. In about 20% of breast cancer cases amplification of the Septin9 gene occurs in double-minute chromosomes, leading to overexpression related to a higher invasion and migration ability of tumor cells [26–28]. Moreover, isoform-specific differences were also shown in interactions with proteins of distinct subcellular organelles and functions [29]. The truncated isoforms 4 and 5 interacting with any member of Septin6 group play a role in cellular signaling and protein ubiquitination, whereas isoform 1 mainly interacts with only Septin8 and regulates cytoskeleton rearrangements. The reason for the excess of certain transcript variants is not known, but epigenetic changes are suspected.

In the light of the above, there are 2 things to consider. On one hand, in contrast to normal ovaries and mammary glands, Septin9 expression remains relatively high post embryonically in the colonic epithelium (and within crypt stem cells), although its level changes with age [21]. On the other hand, the previously mentioned characteristic Septin9 hypermethylation is present at such a high rate even in early-stage COADs that it has been successfully used in diagnostics as an early biomarker of colorectal tumors for over a decade [6–10]. However, very little is known about the function and biology of Septin9 itself in COADs: the composition of splice variants and isoforms in the normal and tumor colonic epithelium, and how and what is affected by methylation. Understanding Septin9 function and the regulatory mechanisms may open up new perspectives for entirely new therapeutic strategies of COADs.

## Results

We examined the differentially methylated amplicons identified in [5] using the UCSC Genome Browser [30], and found them to be localized in a CpG island of the promoter of the *SEPTIN9* v2 transcript variant (Supplementary figure 1A). As very little information is available on the exact function of Septin9 in CRC, we used CRISPR screen data on cell lines in the Cancer Dependency Map (DepMap) Portal [31,32] to investigate the impact of knocking out the gene on tumor growth (Supplementary figure 1B). The Septin9 KO had a negative impact on cancer cell line growth.

Next, we defined the expression pattern of Septin9 splice variants using the TSVdb web-tool [33] which encompassed the RNA-seq isoform expression data of the TCGA Database. In general, Septin9 was expressed in a similar level in tumor and normal samples (Figure 1A and B). However, when comparing expression of different splice variants, we found that the v2 isoform had overall higher expression in the normal samples compared to tumor ones. Similar differences were identified in case of the v6 isoform level as well, although the expression was overall much lower in the normal samples (Figure 1A and 1B). Interestingly, colon adenocarcinoma patients with the lowest expression of the v2 isoform (lower 25%) had significantly better overall survival (p=9.3e-03, HR=0.39), which significance has still retained after adjusting for the known clinical-pathological parameters such as MSI status or stage (p=0.034, HR=0.44; Supplementary figure 1C and D).

**Figure 1.**
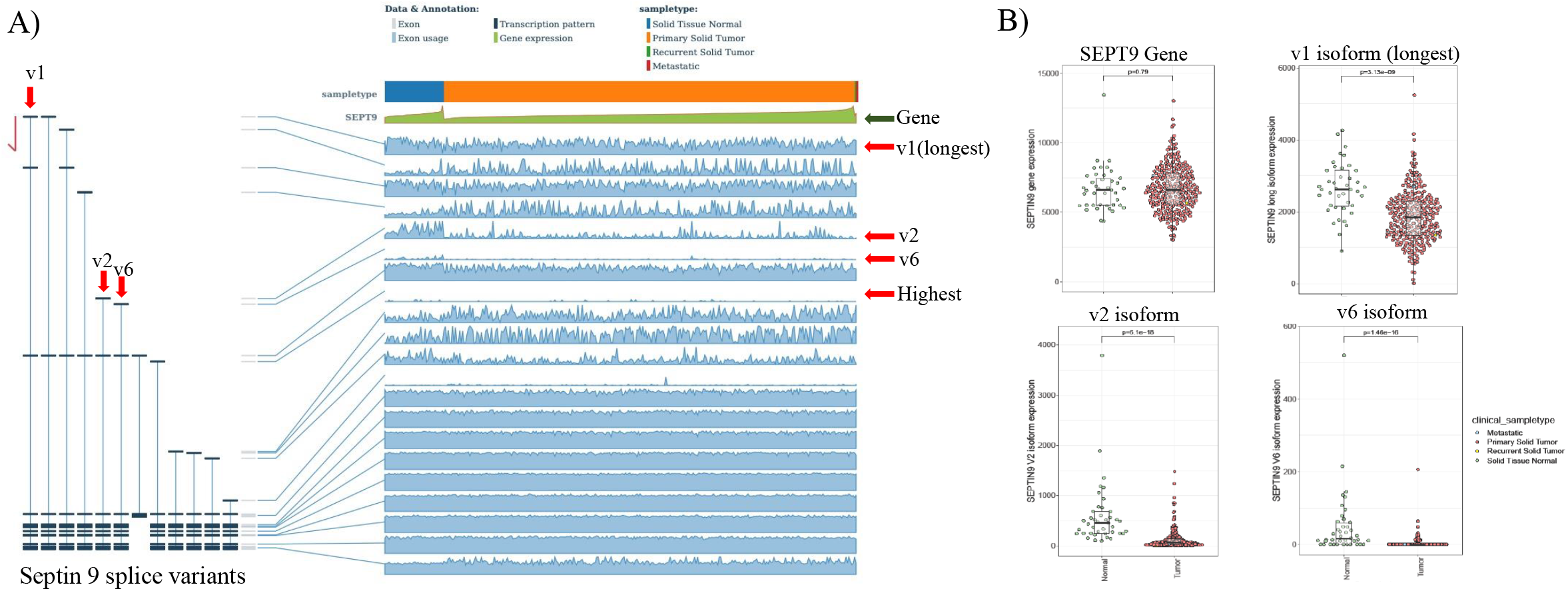
Septin9 splice variant expression analysis using TSVdb tool [33]. (A) Splice variant expression differences were found between tumor and normal samples indicated by red arrows. (B) Septin9 overall expression and mRNA level changes of the longest (v1) and the altered v2 and v6 splice variants in tumor and normal samples.

Chromatin accessibility correlates with genome activity, thus we hypothesized that it may relate with gene expression. Therefore, chromatin accessibility data (normalized signal and quantified peaks from ATAC-seq available in TCGA Database) of the peaks overlapping promoters was compared to the expression level of corresponding isoforms (Figure 2). A moderate but significant positive correlation was found (v1: R=0.56, p=0.0053 in Fig2A-B, v2: R=0.42, p=0.039 in Fig2C-D), where a more open chromatin resulted in higher expression.

**Figure 2.**
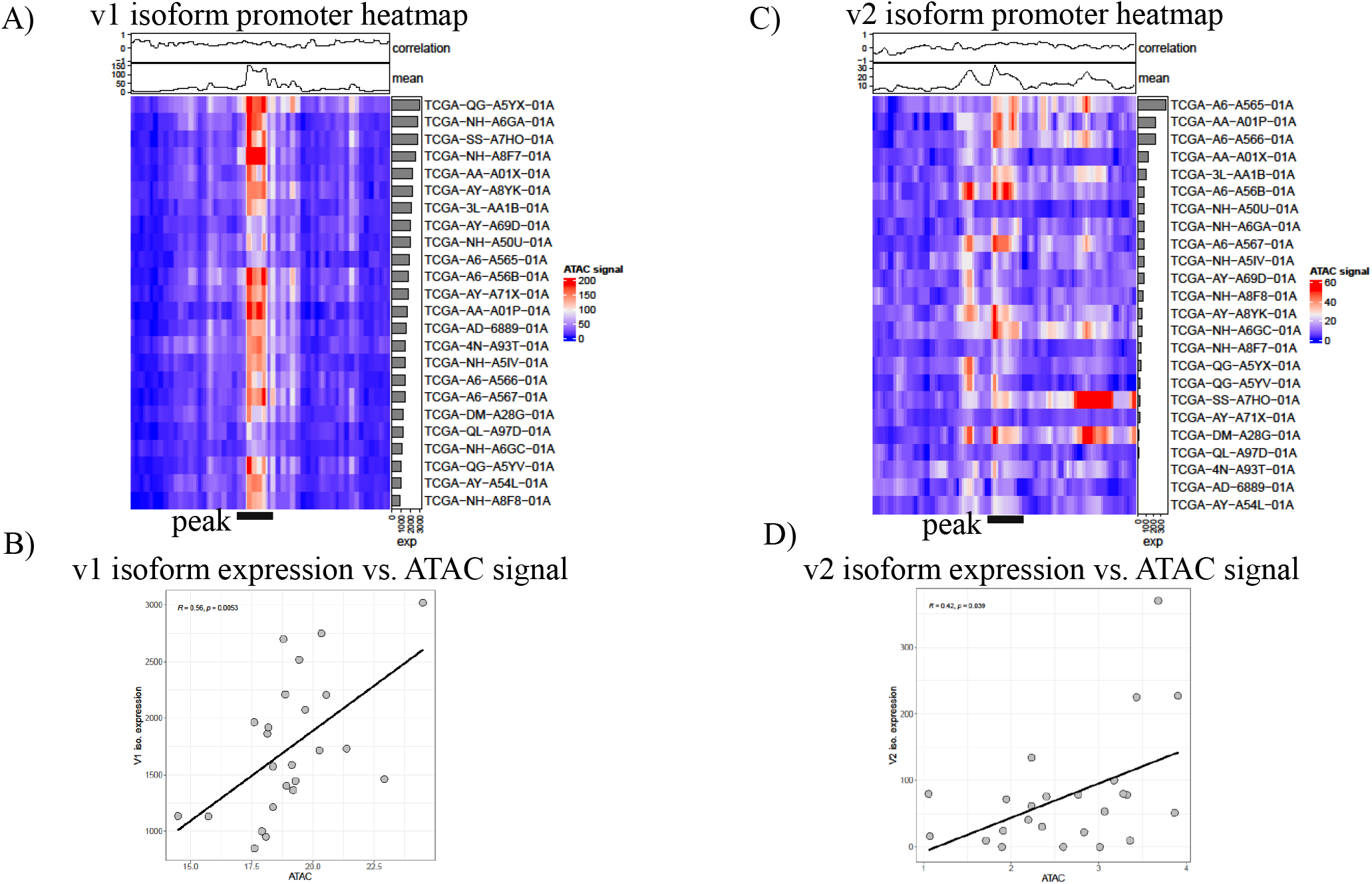
Chromatin accessibility analysis (ATAC-seq) of the longest v1 and altered v2 isoforms available in the TCGA Database. (A) Heatmap of chromatin openness of the Septin9 v1 promoter region. (B) Correlation between v1 variant expression and corresponding ATAC signal. (C) Heatmap of chromatin openness of the Septin9 v2 promoter region. (D) Correlation between v2 variant expression and corresponding ATAC signal.

Besides the *in-silico* analysis of chromatin availability (ATAC-seq), we also examined the signs of chromatin activity using H3K27ac (marker for active enhancers) ChIP-seq data of 10 normal colon and 10 CRC samples available in the TCGA Database. Visualization with IGV (Integrative Genomics Viewer) [34,35] and statistical analysis identified no significant difference in the H3K27 acetylation levels between normal and CRC samples (Supplementary figure 2A-B).

DNA methylation at promoter sites and promoter-CpG islands is a well-known negative regulator of gene expression. We compared the promoter methylation intensity of different Septin9 splice variants in tumor samples to normal ones. Beta values were highly similar in tumor and normal samples in almost all regions (Figure 3A and B). For example, the promoter CpG island of the longest v1 variant is hypomethylated in both normal and tumor samples, and no significant correlation was found with gene expression (Figure 3C and D, Supplementary figure 3A). The only exception was the promoter CpG island of v2 variant (Figure 3A and B) where tumor samples have a significantly higher overall level of methylation. Increased CpG methylation level resulted in overall decreased expression of the v2 isoform (Figure 3E and F, Supplementary figure 3A). Methylation analysis of cell lines from the GDSC Database also found negative correlation of the v2 isoform promoter CpG methylation and expression (Supplementary figure 3B and 3C).

**Figure 3.**
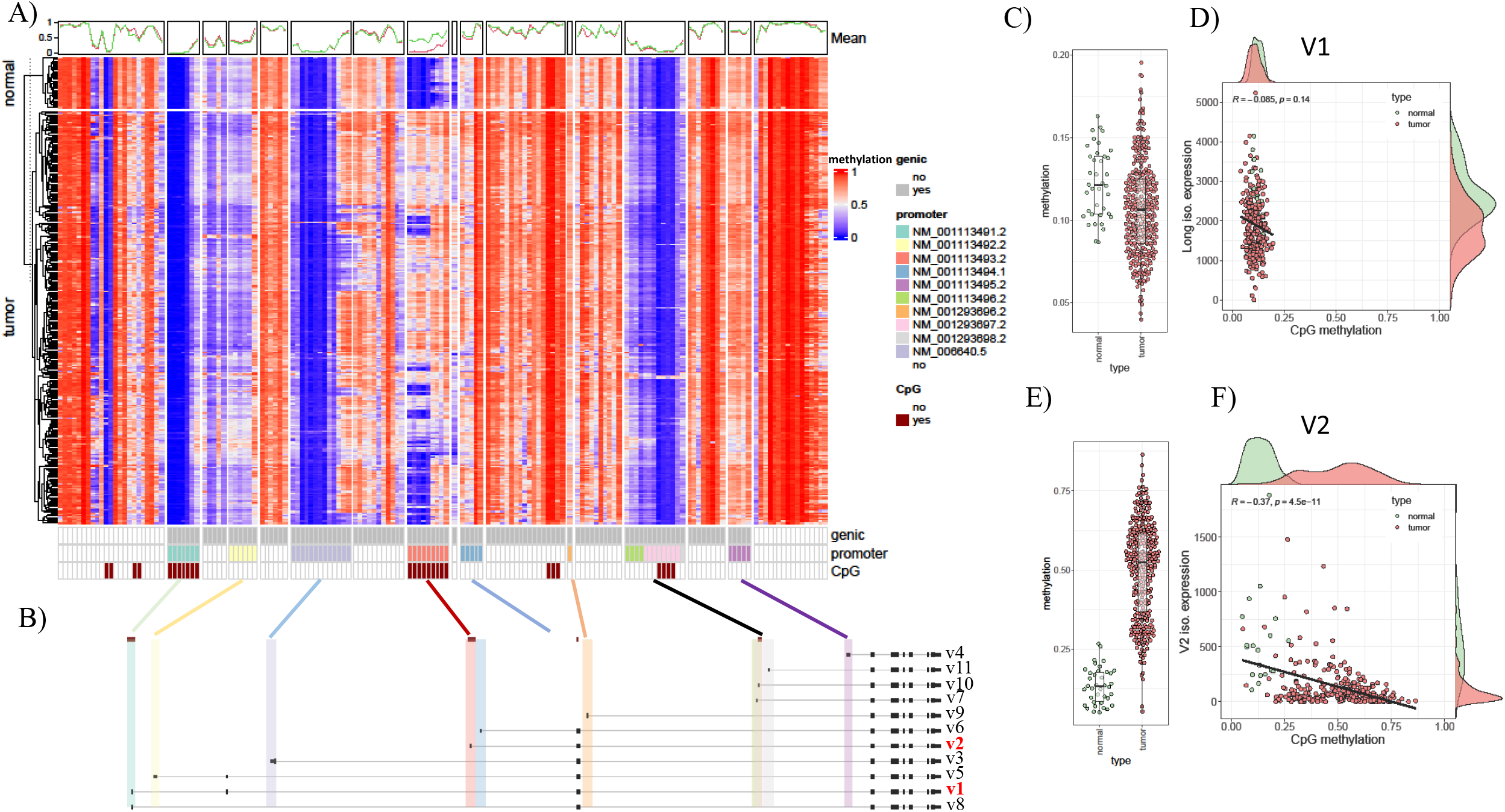
Relationship of promoter methylation intensity of distinct Septin9 variants and their mRNA expression based on data available in the TCGA Database. (A) Heatmap of Septin9 methylation levels in tumor and normal samples. (B) Highlighting the promoter region of different splice variants. (C) Septin9 v1 promoter CpG methylation in tumor and normal samples, and (D) correlation of the longest variant methylation with gene expression. (E) Septin9 v2 promoter CpG methylation in tumor and normal samples, and (F) correlation of the altered variant methylation and gene expression.

Based on our preliminary results using relative quantification by variant specific qRT-PCR, v2 and v6 were also highly underrepresented in COAD cell lines (SW620, HT29 and HT115) comparing to normal colon biopsies (N=4). However, no remarkable differences were observed in the expression of individual variants between the different normal samples (Figure 4A).

**Figure 4.**
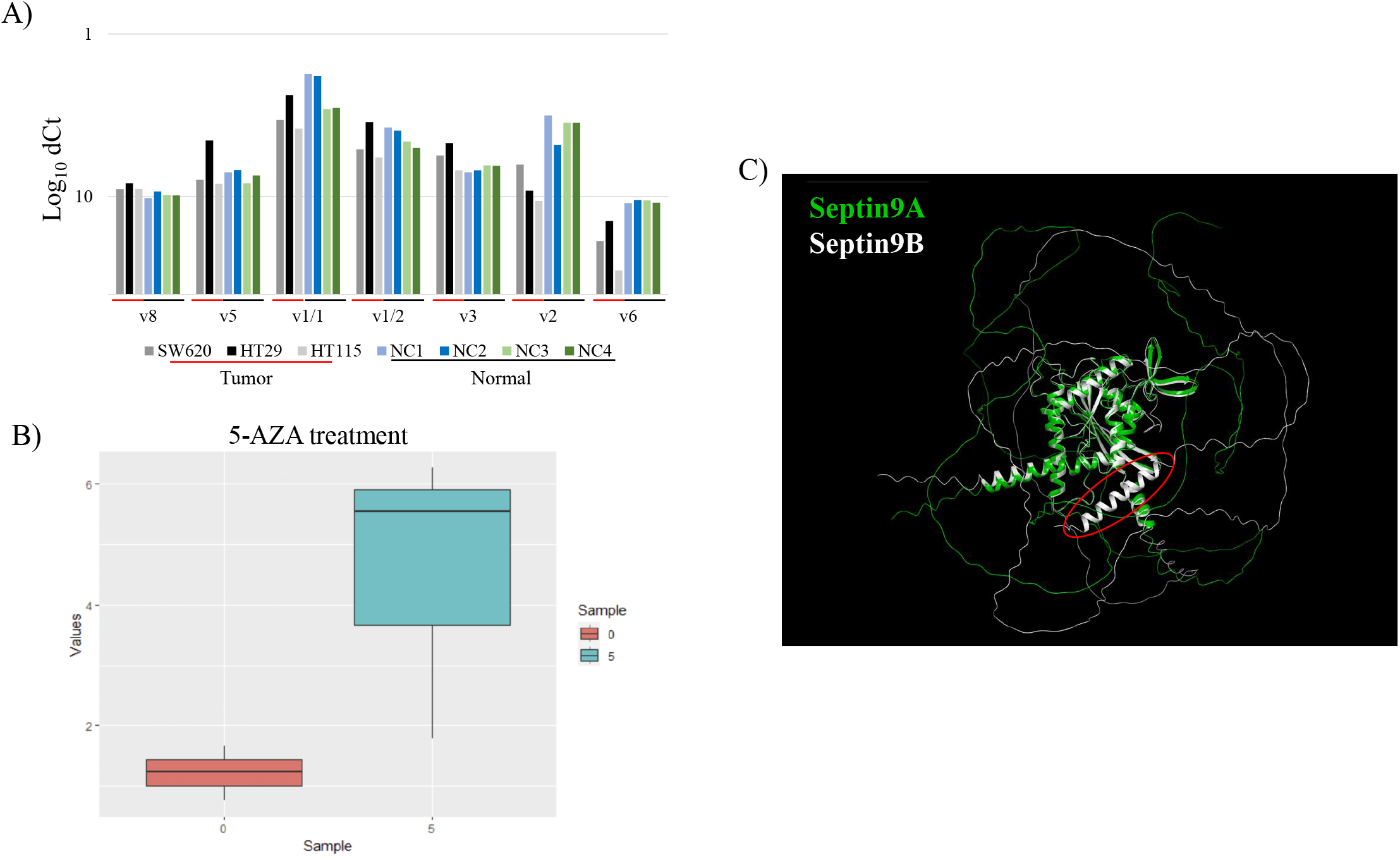
(A) Variant specific qRT-PCR analysis of colon adenocarcinoma cell lines (N=3) and normal colon biopsies (N=4). Relative quantification was performed with Livak method using GAPDH, ACTB, HPRT1 and ALAS1 as internal control genes. (B) 5-AZA treatment (5 μM) effect on the mRNA expression of Septin9 v2 isoform. Boxplots show the average value of triplicates. P-value more than 0.05 was considered statistically significant using Mann-Whitney-Wilcoxon test. (C) Structural illustration and differences of protein isoform ‘A’ (coded by v1) and isoform ‘B’ (coded by v2) using the AlphaFold2 Database [57,58].

In-silico analysis of 5-AZA treatment of 44 COAD cell lines revealed that hypermethylated region of Septin9 is slightly, although not significantly, affected by the global demethylation agent (Figure 4B, Supplementary figure 4A).

It seems that v1 (coding ‘A’ protein isoform) and v2 (coding ‘B’ protein isoform) are the main variants in normal colon, and a decrease in v2 expression due to promoter hypermethylation may have some benefit for the tumor. To reveal possible functional differences between the longest v1 and the altered v2 variants, protein structures were depicted using AlphaFold2. The ‘B’ isoform carries an extra helical structure at the N-terminal end of the protein besides some disordered region missing completely in the ‘A’ isoform (Figure 4C, Supplementary figure 4B).

## Discussion

The expression of Septin9 has been shown to have both tumor suppressor and promoting effects in other cancers, but its complete absence can be lethal to cells [16,23–29]. Hypermethylation of a specific region of the Septin9 gene at the v2 promoter region is a well-known early event in CRC formation [5]. As promoter methylation at CpG sites is a common negative regulator of gene expression, we examined both the overall and splice variant specific expression differences of Septin9 between normal and tumor samples. Our variant-specific expression quantification (Figure 4A), in agreement with the results of *in-silico* analysis of TCGA-COAD RNA-seq data, revealed that the v2 variant is the major target of DNA methylation which could benefit the tumor and can be used as a circulating DNA biomarker for early cancer detection [36]. The level of the v6 variant was also reduced during normal to tumor transition, but its importance may be minor as its expression was already low in normal samples, unlike in breast and ovary carcinomas where a shift from the v6 to v5 variant is frequently observed [16,24,25]. In normal colon samples, the v1 variant was found to be the dominant variant in addition to v2, whose expression was not affected by silencing. In general, global Septin9 expression is retained in COAD, but the normal splice variant composition is altered as an early event in the development of CRC. Gene expression is a multi-level regulated process in which the presence of active DNA is of primary importance [37,38]. To understand the processes underlying the silencing of Septin9 v2, we also examined chromatin accessibility using TCGA ATAC-seq data and H3K27ac ChIP-seq data a marker for active promoters and enhancers from human tissue samples. We observed moderate correlation between ATAC-seq and RNA-seq data for v2, as compared to the correlation for v1. Our analysis of H3K27ac ChIP-seq data did not reveal any significant differences in H3K27ac signals at the v2 promoter region between normal and tumor samples. These findings suggest that promoter methylation of v2 may affect chromatin accessibility, but does not seem to significantly influence H3K27ac signals.

However, a clear and strong association between promoter CpG methylation and v2 expression was observed in both TCGA tumor tissues and GDSC cell line data, with methylation having an effect almost exclusively on the v2 variant. Therefore, unlike breast cancer where v2 is a non-dominant variant [27], it would be necessary to understand the basic role of this isoform in CRC pathogenesis also in relation with the expression of the dominant v1. Assuming that demethylation of the Septin9 v2 promoter may recover its function, we analyzed the effect of global DNA demethylation by investigating a 5-Aza-2’-deoxycytidine treated HT-29 CRC cell line publicly available RNA-seq data. We found evidence of gene expression increase at the methylated Septin9 region, suggesting that a targeted epigenetic modification should be designed for further experimental works.

Previous studies showed that Septin9 interacts with a wide range of proteins, mainly through its C-terminal region [18,39–41], but some proteins, such as FLNA and SH3KBP1, interact through the N-terminal region [18,41], affecting cytoskeletal organization and vesicle transport, among others, in relation to apoptosis. Since the suppression of v2 presumably confers advantage to the tumor, while the other dominant variant, v1 is virtually intact. Therefore, we examined differences in protein structure and hypothesize that aberrant isoform composition may impact interacting partners. We found minor difference in length comparing the v1 and v2 amino acid sequence, correspond to Septin9 variant ‘a’ and ‘b’, respectively. These two isoforms have identical C terminus, but relatively different N terminus amino acid composition. Alterations in length and structure of N-terminal protein region may even cause functionally distinct protein isoforms [20]. Our AlphaFold2 analysis revealed an extra helical structure in the ‘B’ isoform providing it some possible specific function. It previously observed that isoform composition determines the higher-order organisation of filaments [20,25]. Domination of the longest ‘A’ isoform in septin octamers is resulted in a more stable binding of nonbundled microtubules. Therefore, cells may be more sensitive for the treatment with microtubule depolymerization agents such as nocodazole [20]. According to these findings, we hypothesize that aberrant isoform composition may impact on the interacting partners. In conclusion, based on the above information, functional studies are needed to investigate the possible role of v2 variant silencing in colorectal cancer.

## Materials and methods

### Bioinformatic analysis

#### Code availability

All software and bioinformatic tools used in the present study are publicly available.

#### TSVdb

Isoform and gene expression data of SEPTIN9 of the TCGA-COAD dataset was obtained from the TSVdb website [33]. In the search, we selected the SEPT9 gene, summarized by exon. The output contained the SEPTIN9 gene level and isoform level expression, as well as clinical characteristics for the patients used for survival analysis.

#### TCGA methylation

TCGA methylation data for the TCGA-COAD dataset was obtained from the UCSC Xena browser [42] in a sample/probe matrix format. The probes were mapped to genomic coordinates, which were processed and visualized using the *rtracklayer* R package [43]. CpG islands were quantified for each patient by calculated the mean methylation value of each overlapping probe. Methylation heatmap summary was prepared using the *ComplexHeatmap* R package [44].

#### Cell line methylation

Cell line methylation data for was obtained from the GEO under accession number GSE68379 [45]. Raw idat files were processed using the *minfi* package (24478339), and converted to bigwig formats [43,46]. CpG islands were quantified for each patient by calculated the mean methylation value of each overlapping probe. Methylation heatmap summary was prepared using the *ComplexHeatmap* R package [44].

#### Cell line RNA-seq

Cell line RNA-seq data for was obtained from the GEO under accession number GSE68379 [45]. Raw fastq files were trimmed using *trimmomatic* [47], followed by alignment to the hg38 genome using *hisat2* [48]. Raw read counts were calculated using the *TPMcalculator* tool [49]. Normalization scaling factors were obtained from the raw read counts using *DESeq2* [50]. The scaling factors were then used to generate normalized RNA-seq bigwig files using the *BAMscale* tool [51]. Normalized exon-level expression was obtained using the normalized bigwigs and the *rtracklayer* R package [43].

#### ATAC-seq

The normalized COAD ATAC-seq peaks and ATAC-seq signal for the TCGA was obtained from (https://gdc.cancer.gov/about-data/publications/ATACseq-AWG). Analysis of the normalized signal and peak overlapping were performed using the rtracklayer package, followed by heatmap generation using the ComplexHeatmaps

#### H3K27ac ChIP-seq data

Publicly available H3K27ac ChIP-Seq data (GSE166254) in fastq format was downloaded using *SRA-Toolkit*. The fastq files were trimmed using *Trimmomatic*. The overall quality control metrics of fastq were determined by *FastQC*. The trimmed reads were aligned to the hg38 reference genome using *bowtie2. SAMtools* was used for deduplication [52], sorting and indexing. The *MACS2* was applied for peak calling [53]. The bigwig files were created by *BAMscale*. IGV was applied to visualize the bigwig files, specifically for the Septin9 gene (chr17:77279499-77502593), BCAT1 gene (chr12:24808024-24951101), and VIM gene (chr10:17226241-17239593).

#### 5-azacytidine treatment RNA-seq

Publicly available 5-azacytidine treated RNA-seq [54] raw sequence data was downloaded in fastq format by using GEO accession number (GSE41586) using *SRA-Toolkit*. The fastq files were trimmed applying *Trimmomatic* and the overall quality control metrics of fastq files were obtained by *FastQC*. The trimmed reads were aligned to hg38 reference genome using *STAR* aligner [55]. *SAMtools* was used for sorting and indexing of the bam files. The raw read counts were created with the R package *Rsubread* [56]. *DESeq2* was used to determine the scalefactors [50]. *BAMscale* was applied to create the bigwig files, using the previously determined scalefactors. The coverage value of Septin9 exon (chr17:77373341-77373552) was determined with *rtracklayer* R package using the exon coordinates and bigwig files [43]. The bigwigs were visualized using IGV, specifically for the Septin9 gene (chr17:77279499-77502593), Septin9 exon (chr17:77371695-77378018), BCAT1 gene (chr12:24808024-24951101), and VIM gene (chr10:17226241-17239593).

The boxplot was created with the R package ggplot2 using the exon coverage values. Mann–Whitney U test was used to compare the control and 5-AZA treated samples applying the coverage values.

coding: NC_000017.11:77373489-77373557
exon: NC_000017.11:77373336-77373554
chr17:77373341-77373552

#### AlphaFold2 protein structure analysis and visualization

Full-length Septin9A (NP_001106963.1) and 9B (NP_001106965.1) protein models were generated by AlphaFold2. The best ranked models were selected for comparison. The models were aligned with the Schrödinger Protein Structure Alignment module. The amino acid sequence alignment was created with the Schrödinger Multiple Sequence Viewer using aligned structure superposition. Schrödinger, N. Y. N., LLC. Maestro. [57,58]

#### Cell lines and tissue samples

Human colorectal cancer cell lines (SW620, HT29 and HT115) were maintained in DMEM medium supplemented with 10% foetal bovine serum, and 1%-1% Na-pyruvate, hepes, non-essential amino acids and Antibiotic-Antimycotic solution in a 5% CO2 thermostat at 37°C. All chemicals were obtained from Gibco (ThermoFisher Scientific Inc., Budapest, Hungary).

Fresh normal colon biopsies (N=4) were obtained from the 1st Department of Medicine, Semmelweis University (Budapest, Hungary). The study was conducted in accordance with the Declaration of Helsinki and approved by the Ethical Committee of the Medical Research Council of Hungary (Nr.: TUKEB 2005/037).

#### Total RNA extraction and quality control

RNeasy Plus Mini Kit (Qiagen GmbH, Hilden, Germany) was used to isolate the total RNA from cell lines and tissue specimens according to the manufacturer’s protocol. The obtained RNA was quantified using a NanoDrop ND-1000 UV–vis Spectrophotometer (NanoDrop Technologies, Wilmington, Delaware, USA). RNA samples with 260/280 and 260/230 ratios of greater than 1.8 were included in the analysis.

#### qRT-PCR workflow and analysis

Septin9 v1, v2, v3, v5, v6 and v8 variant specific primers, 4 housekeeping genes (GAPDH, ACTB, HPRT1 and ALAS1) and HuNg2 for checking possible DNA contamination were designed by the web-based PrimerQuest tool of the Integrated DNA Technologies (IDT), and purchased from ThermoFisher Scientific Inc. (Budapest, Hungary). Primer sequences are summarized in Supplementary Table 1. Reverse transcription was carried out with the Applied Biosystems High-Capacity cDNA Reverse Transcription Kit at 42°C 2 hours using 2 μg total RNA. cDNAs were stored at −20°C until use. Applied Biosystems SYBR Green PCR Master Mix was used for qRT-PCR experiments running in duplicate on a QuantStudio 6 Real-Time PCR System with the following thermal profile: (1) activation at 95°C for 10 min, (2) amplification (45 cycles): denaturation at 95°C for 15 sec, annealing at 62°C for 20 sec, extension at 72°C for 15 sec, and (3) cooling at 40°C for 30 sec. Each reaction (20 μl in volume) contained 50 ng cDNA and primers in 0.4 μM final concentrations. All the kits and equipment used were from the ThermoFisher Scientific Inc. (Budapest, Hungary). Raw PCR data were analyzed by the Livak method (2^-ddCt^) using the average expression of 4 housekeeping genes as internal control and pooled normal colon biopsies (N=4) as calibrator samples.

## Author contributions

Study design, concept: LV, LSP, SS

Data analysis: LSP, CK, IC, AG, SS

Experimental work: LV, VT, AS

Write and revise manuscript: LV, VT, LSP, SS

## Figure legends

**Supplementary figure 1**. (A) Chromosomal region detected by Septin9 screening kits. (B) Negative impact of Septin9 knocking-out on tumor growth in CRC cell lines using the CRISPR screen data of DepMap Portal [31,32]. (C) Overall survival of Septin9 v2 low (lower 25% of samples) and high patients. (D) Overall survival of Septin9 v2 low and high patients adjusted for the known clinical-pathological parameters.

**Supplementary figure 2**. Chromatin activity (using H3K27ac marker) differences of the Septin9 focusing on the v2 and v6 variants between normal and tumor samples. (A) ChIP-seq data of 10 normal (blue) and 10 tumor (red) samples downloaded from the TCGA Database. (B) Boxplot of the chromatin activity differences between tumor and normal samples.

**Supplementary figure 3**. CpG methylation of altered v2 variant. (A) Probe-level promoter CpG methylation of v1 and v2 isoforms based on TCGA data. (B) Heatmap of the v2 CpG methylation intensity in cell lines from the GDSC Database. (C) Significant negative correlation between Septin9 v2 promoter methylation and mRNA expression based on GDSC Database derived cell line data.

**Supplementary figure 4.** (A) IGV track of the effect of 5-AZA treatment (5 μM) run in triplicate on Septin9 v2 expression. (B) Protein sequence alignment of Septin9 isoforms A (coded by v1 variant) and B (coded by v2 variant).

