## Supplementary figures and images for "Epigenetic Regulation Explains The Functionality Behind Colon Cancer Specific Biomarker Septin9"

### Supplementary figure 1

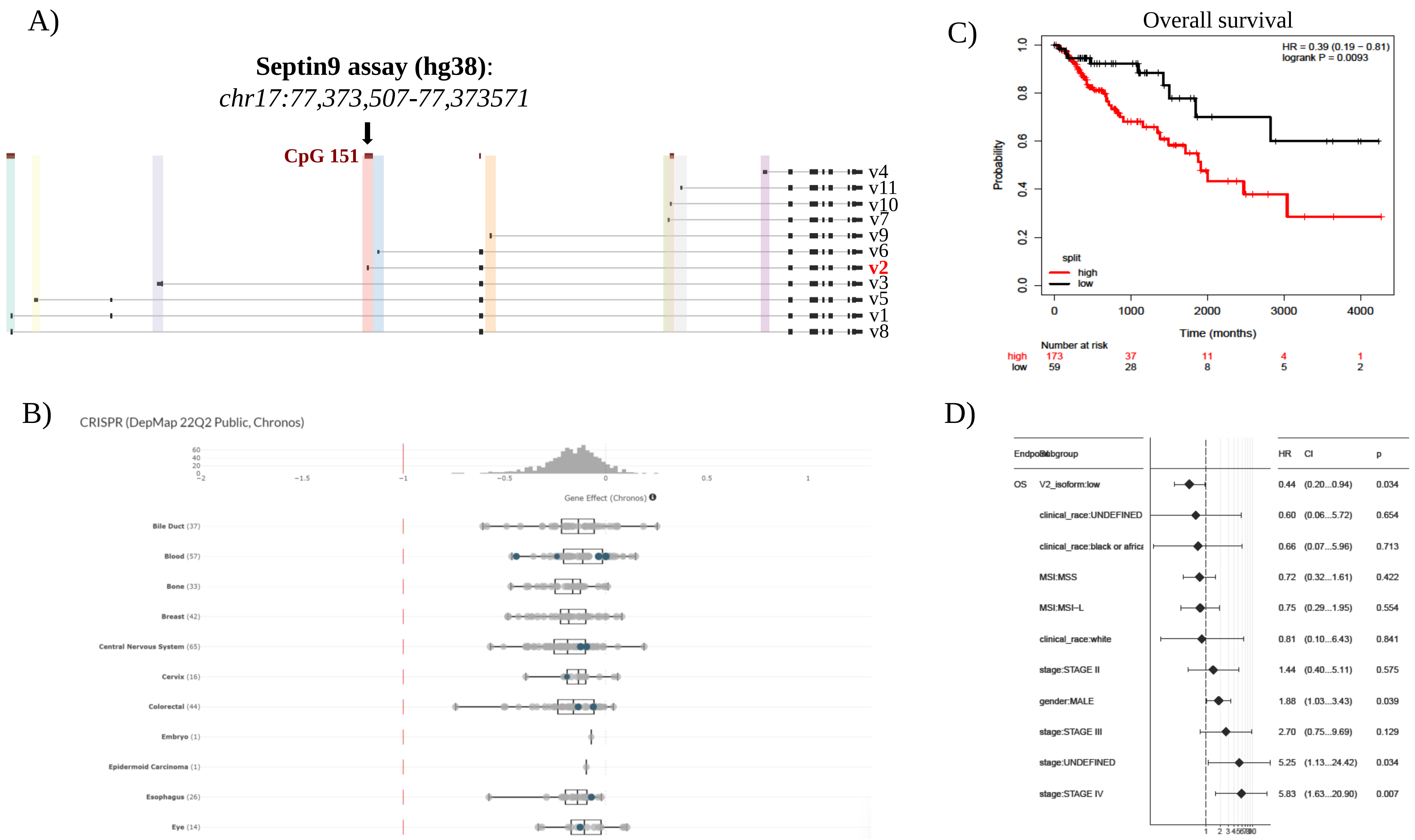

### Supplementary figure 2

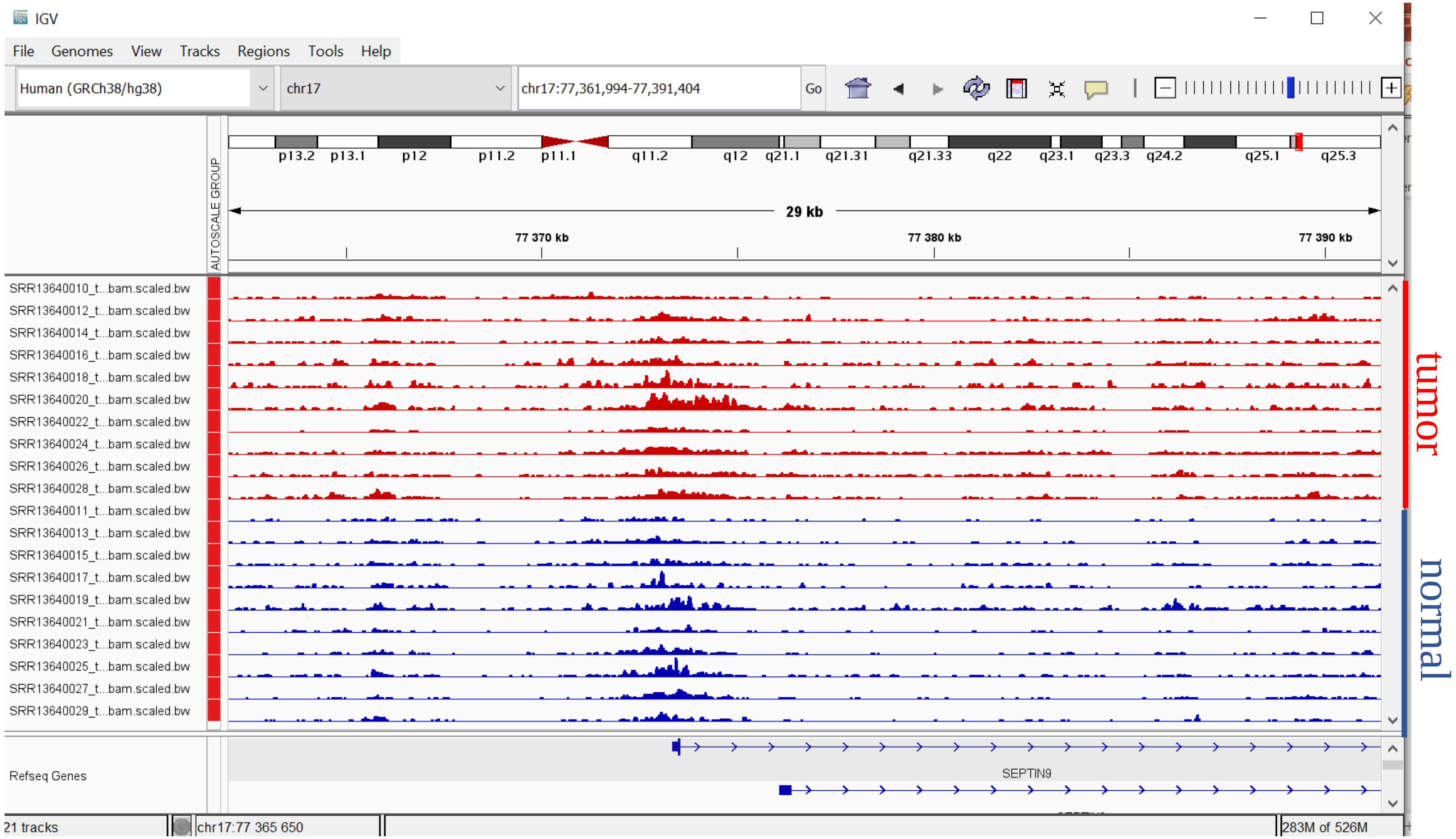

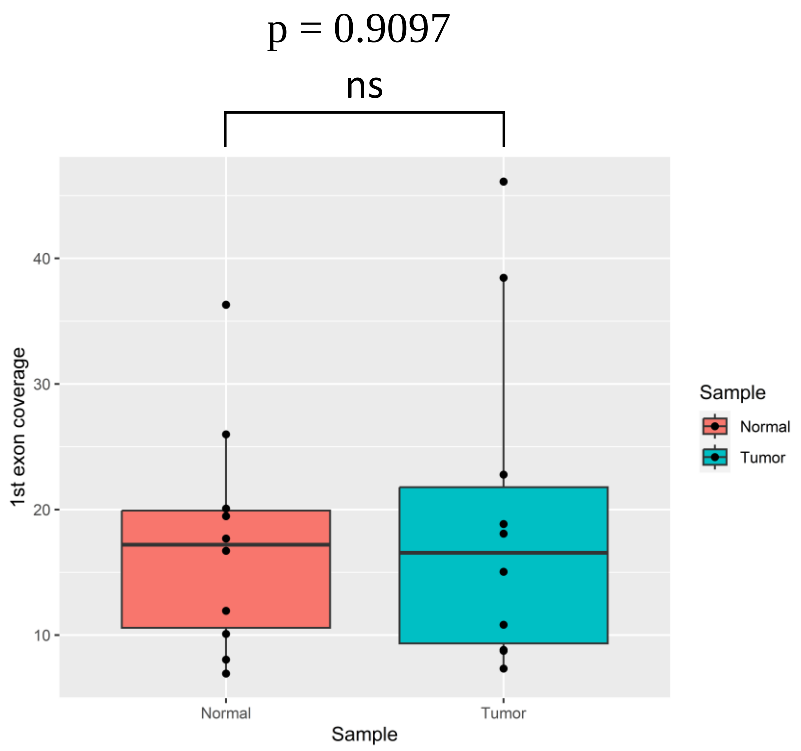

### Supplementary figure 4

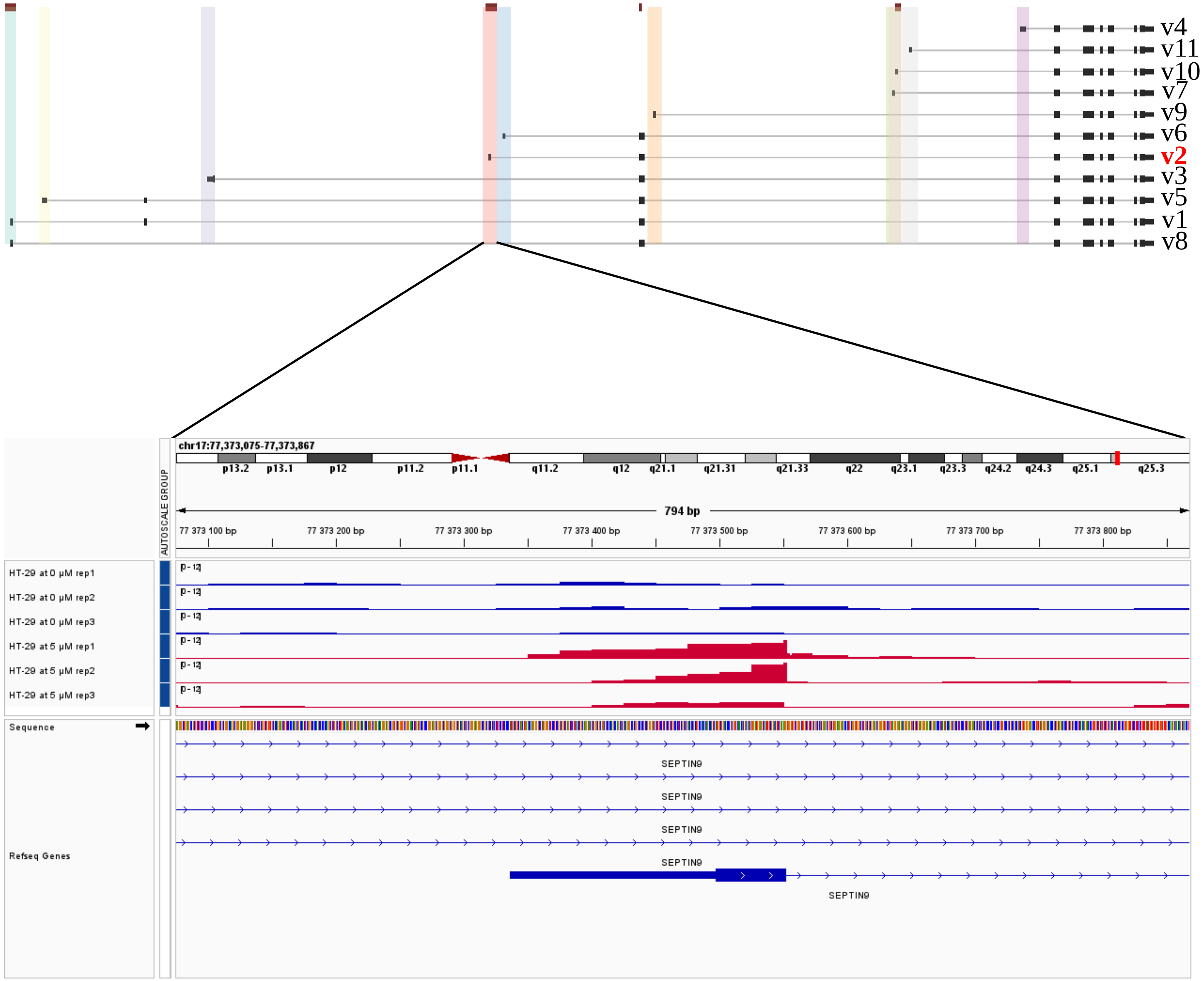

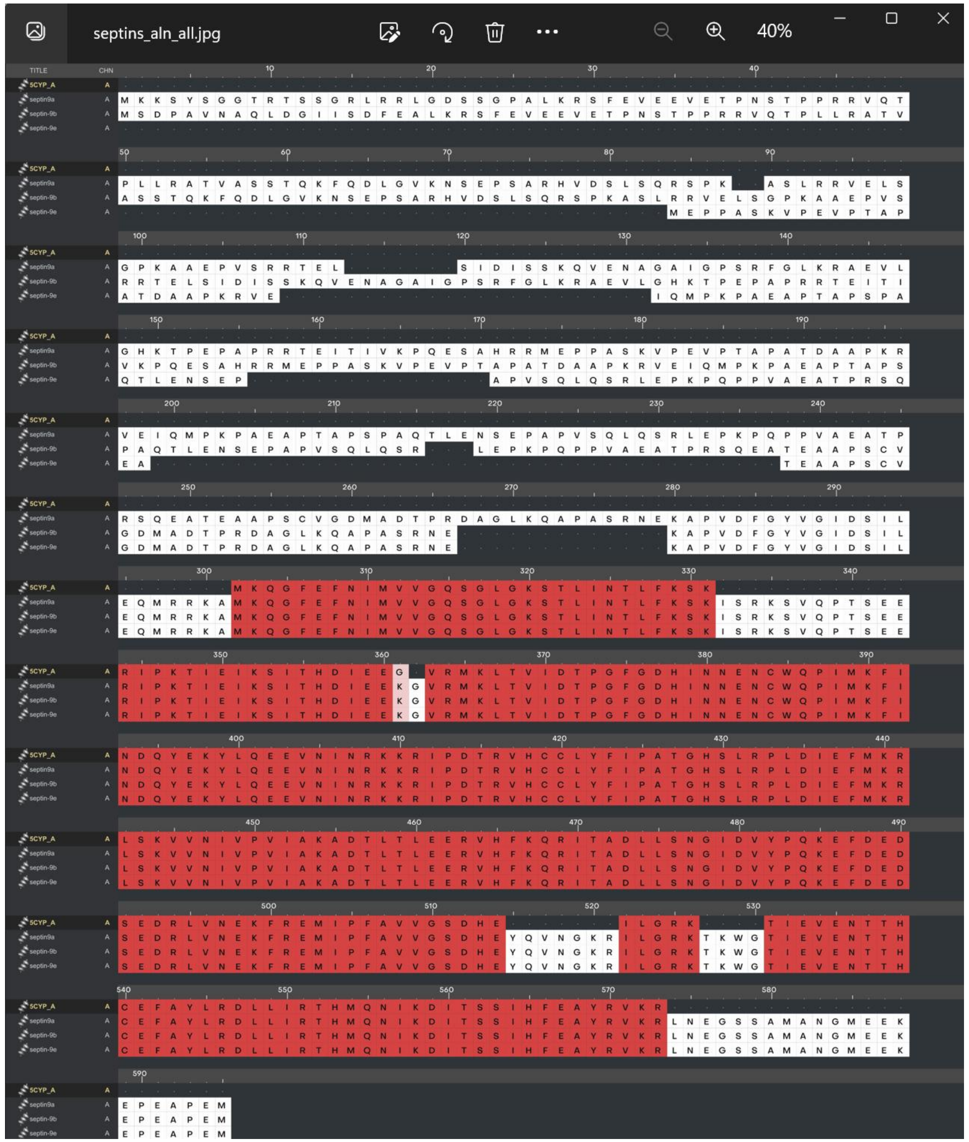
