## Supplementary figure 3 for "Epigenetic Regulation Explains The Functionality Behind Colon Cancer Specific Biomarker Septin9"

A)

### TCGA CpG probe methylation

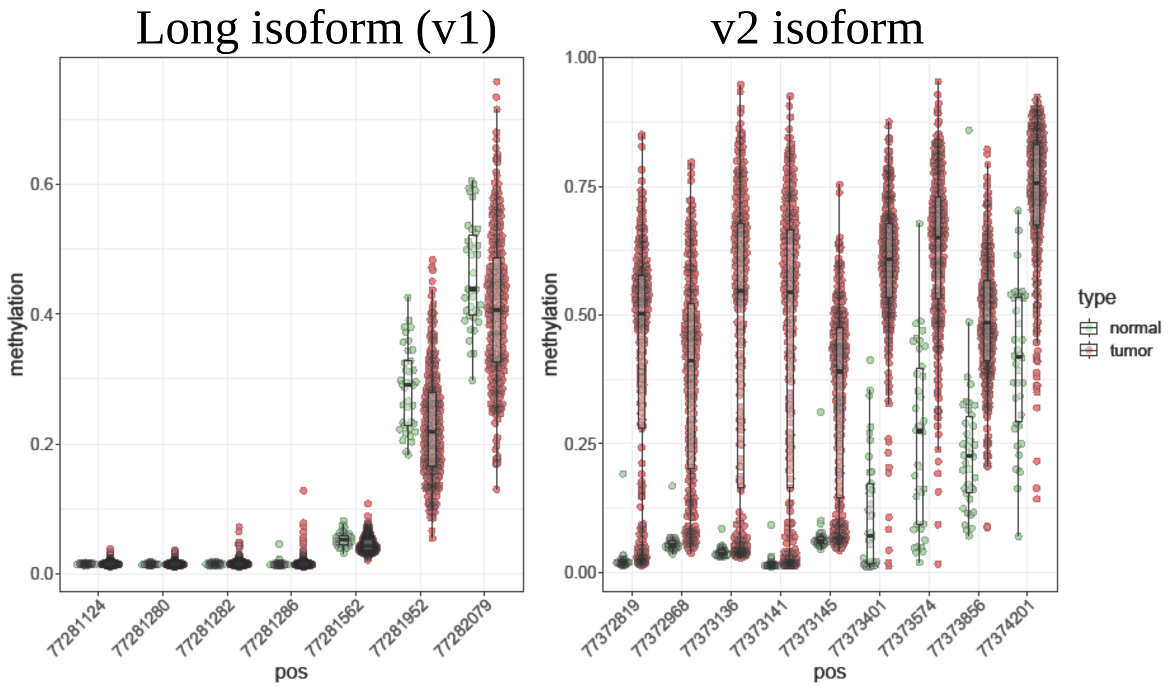


B)

### GDSC (cell line)
### v2 isoform CpG methylation heatmap

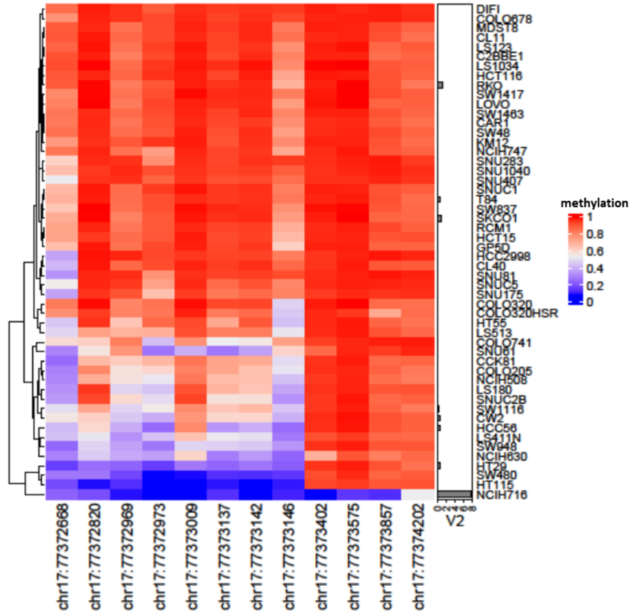


C)

### GDSC (cell line)
### v2 isoform CpG methylation vs. expression

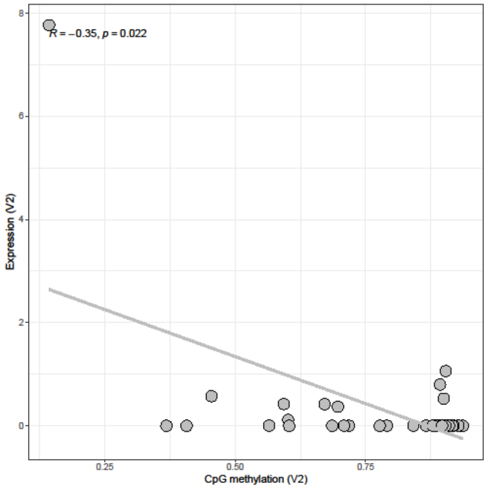
