## Supplementary table 1 for "Epigenetic Regulation Explains The Functionality Behind Colon Cancer Specific Biomarker Septin9"

**Supplementary Table 1.** Primer sequences used in the qRT-PCR experiment.

| Variant | RefSeq | Forward primer (5'-3') | Reverse primer (5'-3') |
|---|---|---|---|
| v1/1 | NM_001113491.2 | AGAAGTCTTACTCAGGAGGCA | AGTTGGGTGTCTCGACCT |
| v1/2 | NM_001113491.2 | GGAGGCACCATGAAGAAGT | GCCACTGGAGTCACCAAG |
| v2 | NM_001113493.2 | GCAGCTGGATGGGATCATT | AGTTGGGTGTCTCGACCT |
| v3 | NM_006640.5 | CCGCTGCTAAATATATCCGTAGG | AGTTGGGTGTCTCGACCT |
| v5 | NM_001113492.2 | GATGTGGCTTCAGTCTCTGTC | AGTTGGGTGTCTCGACCT |
| v6 | NM_001113494.1 | TGGGAGCACAGGTCTCTT | AGTTGGGTGTCTCGACCT |
| v8 | NM_001293695.2 | AGAAGTCTTACTCAGCCTTGAAA | GTTCTTCACGCCCAGGTC |
| GAPDH | NM_002046.7 | CAATGACCCCTTCATTGACC | GACAAGCTTCCCGTTCTCAG |
| ACTB | NM_001101.5 | CAACCGCGAGAAGATGACCC | GAGGCGTACAGGGATAGCAC |
| HPRT1 | NM_000194.3 | TGAGGATTTGGAAAGGGTGT | CATCTCGAGCAAGACGTTCA |
| ALAS1 | NM_000688.6 | AGGAGGATGTGCAGGAAATG | CCTCCATCGGTTTTCACACT |
| HuNg2 | DNA control | CACCACCAGCCTATTTGTTC | TCCTCAACAGGACGAATGTG |
